# Transcription factor Nrf1 regulates proteotoxic stress-induced autophagy

**DOI:** 10.1101/2023.06.29.547114

**Authors:** Hatem Elif Kamber Kaya, Madison A. Ward, Janakiram R. Vangala, Holly A. Byers, Antonio Diaz, Susmita Kaushik, Ana Maria Cuervo, Senthil K. Radhakrishnan

## Abstract

Cells exposed to proteotoxic stress invoke adaptive responses aimed at restoring proteostasis. Our previous studies have established a firm role for the transcription factor Nuclear factor erythroid derived 2-related factor 1 (Nrf1, also called NFE2L1) in responding to proteotoxic stress elicited by inhibition of cellular proteasome. Following proteasome inhibition, Nrf1 mediates new proteasome synthesis, thus enabling the cells to mitigate the proteotoxic stress. Here we report that under similar circumstances, multiple components of the autophagy lysosomal pathway (ALP) are transcriptionally upregulated in an Nrf1-dependent fashion, thus providing the cells with an additional route to cope with proteasome insufficiency. In response to proteasome inhibitors, Nrf1-deficient cells displayed profound defects in invoking autophagy and clearance of aggresomes. This phenomenon was also recapitulated in NGLY1 knockout cells (a model for NGLY1 disease) where Nrf1 is known to be non-functional. Overall, our results significantly expand the role of Nrf1 in shaping the cellular response to proteotoxic stress.

## INTRODUCTION

Effective destruction and recycling of misfolded, defective and aggregated proteins is a key step in proteostasis and is essential for cellular survival (Lindquist and Kelly, 2011). This step is facilitated by the ubiquitin-proteasome system (UPS) and the autophagy-lysosomal pathway (ALP), the two major proteolytic pathways in the cell (Pohl and Dikic, 2019).

The central player in the UPS is the 26S proteasome, a multi-subunit proteolytic complex, which recognizes and degrades ubiquitylated substrates. Within the 26S proteasome, the actual process of protein degradation is achieved in the 20S catalytic core that harbors chymotrypsin-like, trypsin-like, and caspase-like activities. Often, the barrel-shaped 20S core subunit is capped on one or both ends by a 19S regulatory particle that aids in substrate deubiquitylation and unfolding, both of which are essential steps for proteasome-mediated degradation (Rousseau and Bertolotti, 2018). Thus, the UPS is better suited for the recycling of soluble proteins and not aggregates. In contrast, ALP via macroautophagy (hereafter autophagy) can readily degrade protein aggregates and even damaged cellular organelles. This pathway involves substrate sequestration in double-membranous vesicles called autophagosomes. The autophagosomes then fuse with hydrolytic enzyme-containing lysosomes thereby generating autolysosomes wherein the cargo is degraded (Pohl and Dikic, 2019).

The proteasome is regulated at multiple levels including transcription, translation, assembly, and posttranslational modifications (Schmidt and Finley, 2014). Our previous studies assign a central role for the transcription factor Nuclear factor erythroid derived 2-related factor 1 (Nrf1, also called NFE2L1) in regulating the transcription of proteasome genes (Radhakrishnan et al., 2014; Radhakrishnan et al., 2010; Vangala and Radhakrishnan, 2019; Vangala et al., 2016). Nrf1 is co-translationally inserted into the membrane of the endoplasmic reticulum as a precursor p120 and is subject to constant degradation under normal conditions. However, when the cellular proteasome is inhibited, Nrf1 accumulates and is cleaved by the protease DDI2 resulting in p110, an active form of this transcription factor which then migrates to the nucleus and activates proteasome genes (Koizumi et al., 2016; Lehrbach and Ruvkun, 2016). Thus, Nrf1 is designed to sense and respond to proteasome dysfunction, thereby alleviating the resultant proteotoxic stress. Given that proteasome inhibitors such as bortezomib, carfilzomib and ixazomib are now being used in the clinic as cancer therapeutics, it is important to understand these cellular adaptive responses to proteotoxic stress. These cellular adaptations are also relevant in the context of neurodegenerative diseases characterized by accumulation of protein aggregates which are known to impair proteasome function (Myeku et al., 2016).

The UPS and ALP have long been considered as two parallel and independent intracellular degradation pathways. However, recent findings have challenged this notion, wherein autophagy has been shown to act as a compensatory pathway when the proteasome is impaired (Bao et al., 2016; Ding et al., 2007; Zhu et al., 2010). Despite this evolving view, the molecular mechanisms that connect these proteolytic pathways are not completely understood. In this study, we present evidence to demonstrate that Nrf1 could act as a molecular link between these two degradation systems and that it could mobilize autophagy when the proteasome is impaired.

## RESULTS

### Nrf1 upregulates the expression of Autophagy-Lysosomal Pathway (ALP) genes in response to proteasome inhibition

Our previous studies have demonstrated that inhibition of cellular proteasomes mobilizes the Nrf1 pathway resulting in *de novo* synthesis of proteasome genes and subsequent recovery of proteasome activity (Radhakrishnan et al., 2014; Radhakrishnan et al., 2010; Vangala and Radhakrishnan, 2019; Vangala et al., 2016). To gain further insight into other processes that might also be regulated in a Nrf1-dependent fashion after proteotoxic stress, we analyzed our RNA-sequencing (RNA-seq) dataset with NIH-3T3 wild-type and Nrf1-knockout (Nrf1^KO^) mouse fibroblasts exposed to proteasome inhibitor carfilzomib for either 6 or 24 hrs (NCBI GEO accession GSE144817; (Vangala et al., 2020)). As expected, we observed a wide-spread induction of proteasome genes in 6 and 24 hr time points in wild-type but not in Nrf1^KO^ samples (Supplementary Fig S1). Strikingly, we also noticed that a number of autophagy and lysosomal pathway (ALP) genes were upregulated in a Nrf1-dependent manner most prominently in the 24 hr time point, with some genes also showing a milder induction as early as 6 hrs. This observation points to the possible existence of a Nrf1-mediated autophagic response as a result of sustained proteotoxic stress.

**Figure 1.**
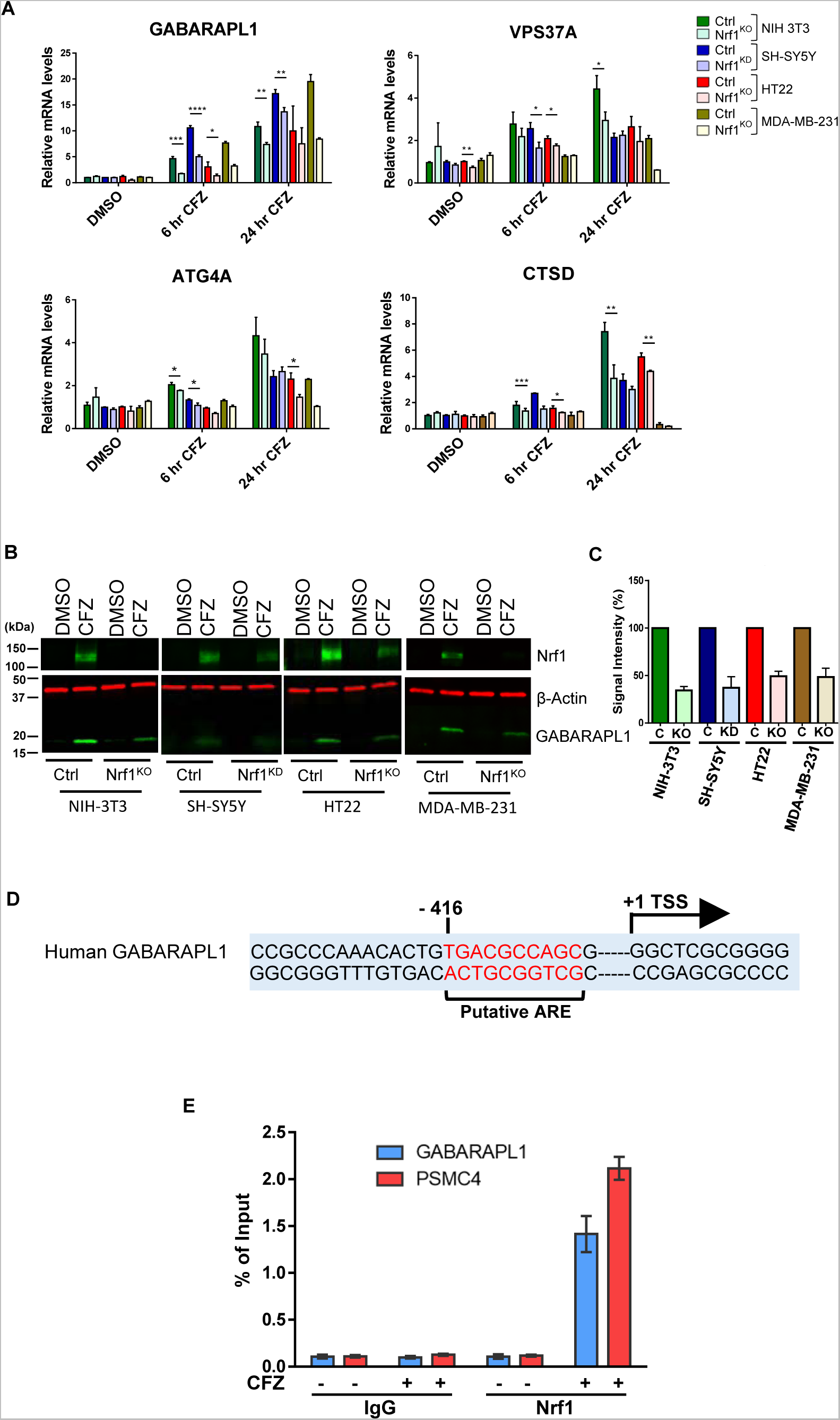
Nrf1 regulates expression of Autophagy-Lysosome Pathway (ALP) genes upon proteasome inhibition. **A.** qRT-PCR analysis of NIH-3T3, SH-SY5Y, HT22 and MDA-MB- 231 cells that are control (Ctrl) or Nrf1-depleted (KO or KD) were treated with either DMSO or 200 nM CFZ for 6 h or 24 h. Expression levels of GABARAPL1, VPS37A, ATG4A and CTSD were analyzed using gene specific primers as shown. 18s rRNA or GAPDH levels were used for normalization. **B.** Western blot analysis of GABARAPL1 protein in NIH-3T3, SH-SY5Y, HT22 and MDA-MB-231 cells treated with either DMSO or 200 nM CFZ for 24 h. β-Actin was used for loading control. **C.** Quantification of GABARAPL1 signal intensity, normalized to β-Actin signal. 3 biological replicates for each cell line were used to perform qRT-PCR and western blotting. p-values were calculated by student’s t test. *<0.5, **,0.05, ***<0.005, ****<0.0005. **D.** Putative ARE sequence close to the transcription start site (+1) of human GABARAPL1. ARE sequence and transcription start site are marked. **E.** Chromatin immunoprecipitation of GABARAPL1 and PSMC4 (proteasome gene; positive control) in DMSO, CFZ (200 nM/ 6 hr) treated SH-SY5Y cells were carried out with IgG, Nrf1 antibodies. qPCR analysis completed with primers flanking the putative ARE sequences (Nrf1 binding) in the promoter region. Nrf1 binding to each gene is expressed as percentage of the input. Error bars denote S.D. (n = 3).

To further evaluate the role of Nrf1 in this response, we used a panel of cell lines of different origin (NIH-3T3 murine fibroblasts, SH-SY5Y human neuroblastoma, HT22 mouse hippocampal, and MDA-MB-231 human breast cancer) that are either control or Nrf1-deficient and assayed for changes in transcript levels of select ALP-related genes in response to treatment with the proteasome inhibitor carfilzomib (CFZ). We observed a time-dependent increase in the transcript levels of these genes in control cells in response to CFZ, but this effect was significantly attenuated in Nrf1-deficient cells (Fig 1A). Consistent with these results, GABARAPL1 protein levels robustly increased in response to CFZ in control cells, but this effect was muted in Nrf1-deficient cells (Fig 1B and 1C). To evaluate if Nrf1 could directly transactivate ALP-related genes, we first analyzed the regulatory regions of human GABARAPL1 gene and found a putative antioxidant response element (ARE; the sequence to which Nrf1 is known to bind (Biswas and Chan, 2010; Wang et al., 2007)) in the proximal promoter region (Fig 1D). Importantly, using chromatin immunoprecipitation (ChIP) experiments, we observed recruitment of Nrf1 to this ARE-containing region in response to CFZ in SH-SY5Y cells (Fig 1E). To further extend these results, we examined the ChiP-seq datasets in the Encyclopedia of DNA Elements (ENCODE) database (Luo et al., 2020) and found evidence for the binding of Nrf1 to the regulatory regions of multiple ALP-related genes (Supplemental Table S1). Together, these findings are consistent with a model in which proteotoxic stress-induced Nrf1 transactivates ALP genes via direct binding to their regulatory regions. To rule out drug-specific effects, we used another proteasome inhibitor, bortezomib (BTZ), and found its effect to be similar to CFZ in robustly activating ALP genes in control when compared to Nrf1-deficient cells (Supplemental Fig S2).

Next, we evaluated if reinstating Nrf1 expression in Nrf1-deficient cells could rescue the activation of ALP genes in response to proteasome inhibition. To this end, we overexpressed the Nrf1 precursor p120 in SH-SY5Y and HT22 Nrf1-deficient cells. Regardless of the proteasome inhibitor used (CFZ or BTZ), we saw that the transcripts of ALP genes were upregulated much more in these Nrf1-rescued cell lines, when compared to the Nrf1-deficient cell lines (Fig 2A and Supplemental Fig S3). We observed a similar effect when we examined GABARAPL1 at the protein level (Fig 2B and 2C).

**Figure 2.**
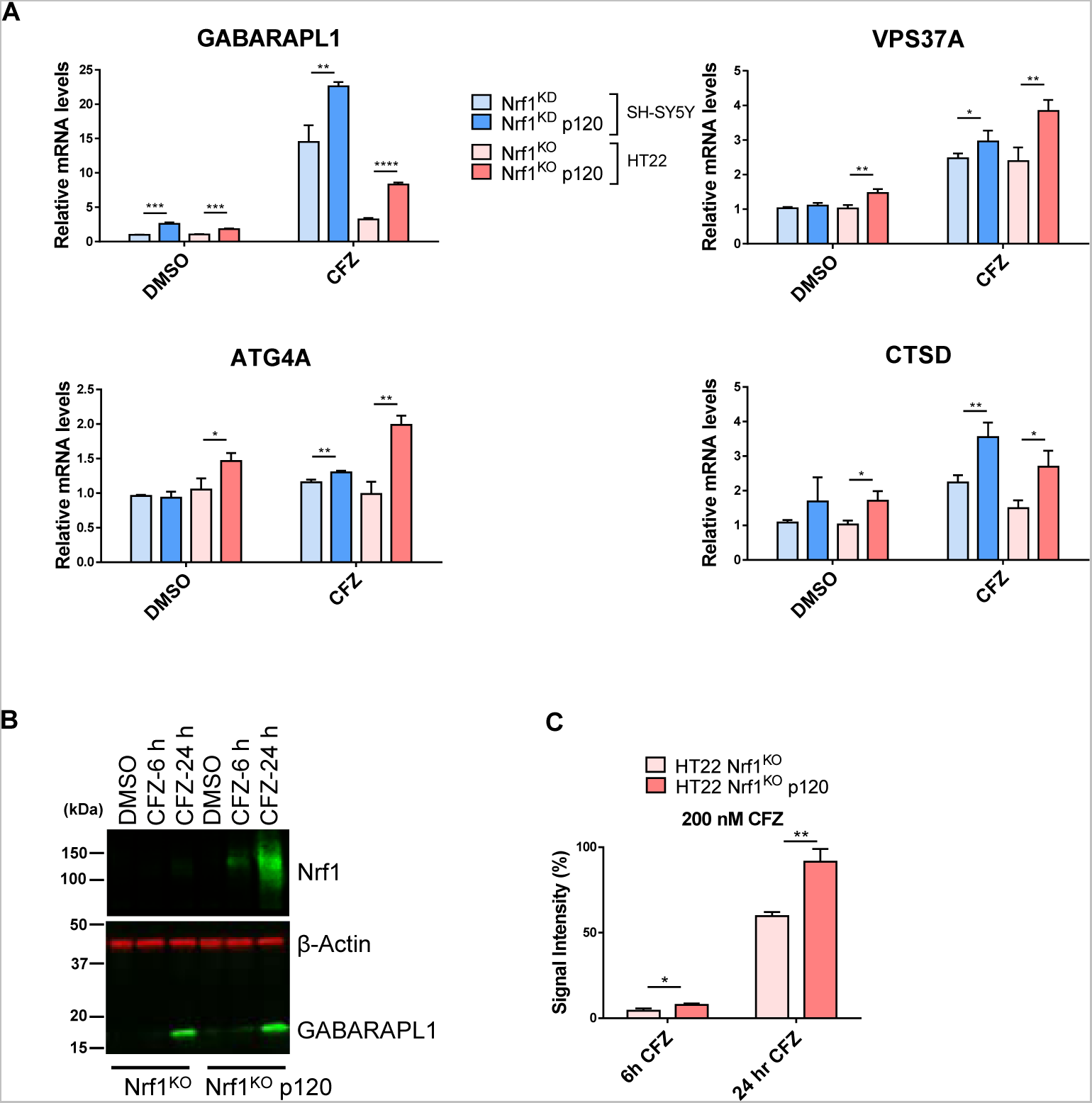
Adding back Nrf1 in Nrf1-deficient cells rescues suppressed expression of ALP genes upon proteasome inhibition. **A.** SH-SY5Y-Nrf1^KD^ and HT22-Nrf1^KO^ cells were infected with Nrf1(p120). Both SH-SY5Y-Nrf1^KD^, p120 rescue and HT22-Nrf1 ^KO^, p120 rescue cells were treated with 200 nM CFZ for 6 h and then analyzed by qRT-PCR to measure the expression levels of indicated genes and mRNA levels of 18s rRNA or GAPDH was used for normalization. **B.** Western blot analysis of GABARAPL1, Nrf1 in HT22-Nrf1^KO^, p120 rescue cells treated with either DMSO or 200 nM CFZ for 6 h and 24 h. β-Actin was used for loading control. **C.** Quantification of GABARAPL1 signal intensity, normalized to β-Actin signal. 3 biological replicates for each cell line were used to perform qRT-PCR and western blotting. p- values were calculated by student’s t test. *<0.5, **,0.05, ***<0.005, ****<0.0005

### Nrf1 induces autophagic flux in response to proteasome inhibition

To understand whether proteasome inhibitor-mediated upregulation of ALP genes by Nrf1 can induce compensatory autophagy, we decided to examine autophagic flux in HT22 cells. LC3 protein is essential for execution of autophagy and is widely used as a marker to assess autophagy activation (Klionsky et al., 2016). We first used a tandem monomeric mCherry-GFP-tagged LC3 protein (N’Diaye et al., 2009) to monitor autophagic flux. GFP is quenched under acidic conditions, while mCherry is stable. Therefore, colocalization of mCherry and GFP signals indicates autophagosomes; whereas just mCherry signal shows autolysosomes due to instability of GFP in lysosomes. In the case of autophagy induction, this reporter yields an increase in both colocalized and red puncta; however, autophagy inhibition causes an increase in colocalized puncta but decrease in red puncta. With the help of this reporter, we observed that autophagic flux is induced in HT22 wild-type cells and inhibited in HT22 Nrf1^KO^ cells in response to BTZ (Fig 3A). We calculated the autophagic flux from three independent experiments by dividing colocalized puncta area with red puncta area and by normalizing this value to total cell number. This calculation suggested that autophagic flux in HT22 Nrf1^KO^ cells are significantly impaired, compared to wild-type cells, upon proteasome inhibition (Fig 3A and 3B).

**Figure 3.**
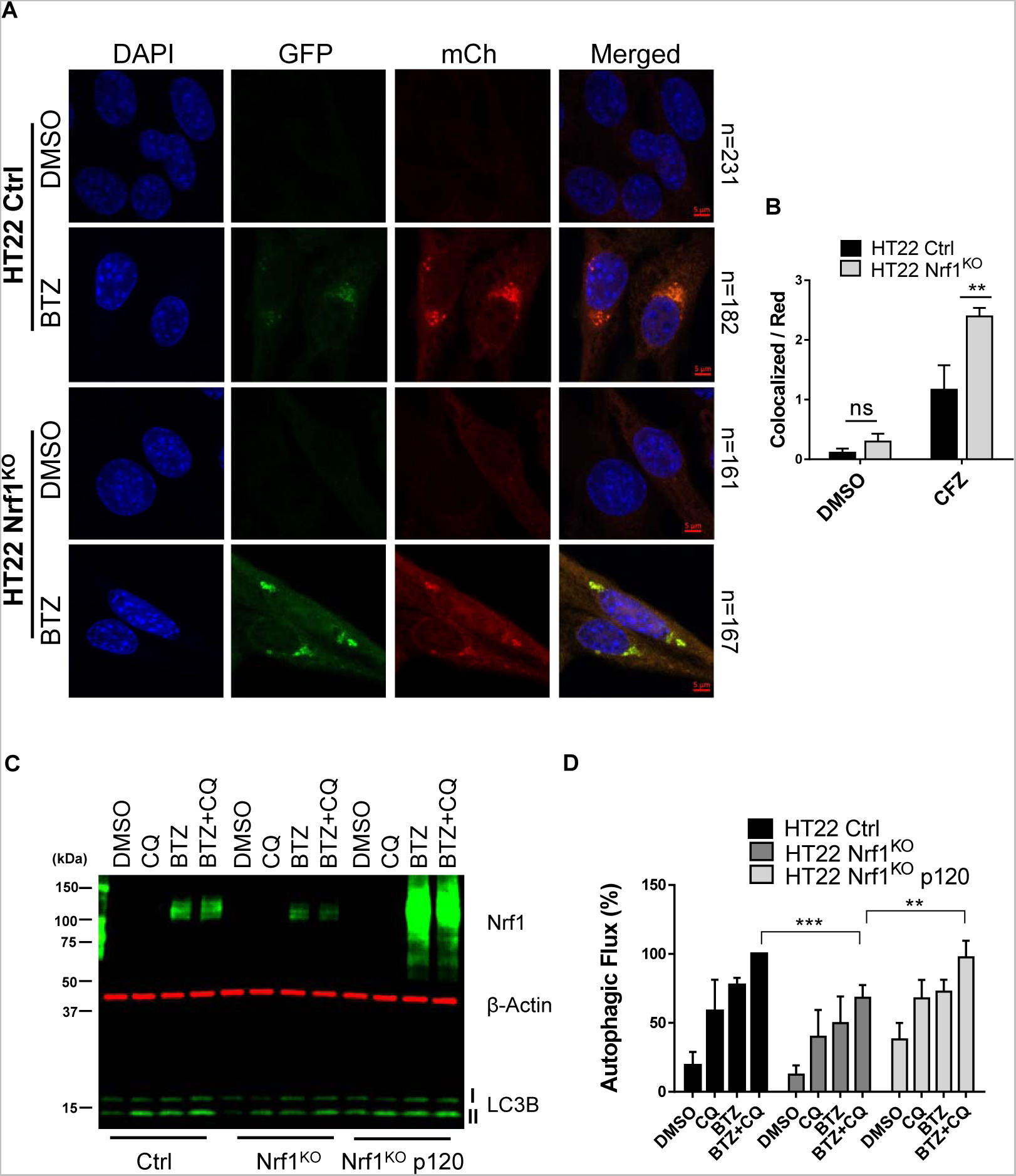
Nrf1 can induce autophagy upon proteasome inhibition. **A.** Confocal microscopy images of HT22 wild-type (control; ctrl) and Nrf1^KO^ cells, stably expressing mCh-GFP-LC3B construct for detecting autophagic flux. Cells were treated either with DMSO or 200 nM BTZ for 20 h. **B.** Autophagic flux is calculated by dividing colocalized puncta signal by red puncta signal and normalizing this level to total cell number in each condition. Number of cells analyzed in each sample is noted next to the images in (A). **C.** HT22-ctrl, Nrf1^KO^, Nrf1^KO^ +p120 cells were treated with DMSO, 200 nM BTZ (20h), 60 μM CQ (3 h) or both BTZ (20 h) and CQ (3 h). Western blot analysis of Nrf1 and LC3B completed with specific antibodies. β-Actin was used as a loading control. **D.** Autophagic flux is calculated by normalizing LC3B-II levels to β-Actin levels from C. 3 biological replicates for both microscopy and western blotting were used. p-values were calculated by student’s t test. *<0.5, **,0.05, ***<0.005

Next, we wanted to validate our observations via an orthogonal setup. Autophagic flux can be tested by using an autophagy inhibitor with or without the potential autophagy inducer and then examining LC3-II turnover by western blot (Klionsky et al., 2016). Specifically, pro- LC3 protein is first cleaved by ATG4 to form LC3-I which is then lipidated to generate LC3-II. This lipidated LC3-II then conjugates to autophagosomes and recruits autophagic cargo by interacting with autophagy receptors such as p62 (Melia et al., 2020). Interestingly, both inhibition and activation of autophagy can result in elevated LC3-II level due to inefficient degradation of LC3-II and increased production of LC3-II, respectively (Mizushima and Murphy, 2020). Accordingly, we used a lysosomal inhibitor, Chloroquine (CQ) to inhibit and BTZ to induce autophagy. Wild-type HT22 cells showed an increase in LC3-II level, when treated with either CQ or BTZ, compared to DMSO only treatment (Fig 3C). Moreover, when we treated wild-type HT22 cells with both CQ and BTZ, we observed an even higher increase in LC3-II level, compared to single treatments, confirming the role of BTZ in inducing autophagy (Fig 3C). In contrast, we observed that LC3-II protein level was significantly less in Nrf1^KO^ cells compared to wild-type cells in response to both CQ and BTZ (Fig 3C and 3D), suggesting an impairment of autophagic flux in Nrf1^KO^ cells. Next, we examined whether restoring Nrf1 levels in Nrf1^KO^ cells can rescue the autophagic flux impairment. We observed that overexpression of precursor Nrf1 p120 in Nrf1^KO^ cells significantly increased LC3-II levels, compared to Nrf1^KO^ cells, when treated with both CQ and BTZ (Fig 3C and 3D). Overall, our data suggest that Nrf1 is required for activation of compensatory autophagy upon proteasome inhibition. Consistent with this theme, we also found that NIH-3T3 Nrf1^KO^ cells display defects in basal and starvation-induced autophagy (Supplemental Fig S4).

### Nrf1 is required to clear aggresomes associated with proteasome inhibition

Functional decline in the activity of the proteasomes through aging, genetic mutations or environmental stress can lead to aggregation of misfolded and unwanted proteins (Lopez-Otin et al., 2013). These toxic aggregates such as in the case of neurological diseases are also known to inhibit proteasomes (Myeku et al., 2016). These aggregates form perinuclear inclusion bodies that are called aggresomes, and can be cleared by a selective type of autophagy termed as Aggrephagy (Bauer et al., 2023). To test whether Nrf1 plays a role in the aggresome clearance pathway, we treated HT22 wild-type and Nrf1^KO^ cells with CFZ and observed them under the confocal microscope. Treatment of these cells with CFZ induced aggresome formation, which was evident by staining these cells with a polyubiquitin antibody FK2 (Fig 4A). We then removed CFZ and let the cells recover in fresh complete media and observed for residual aggresomes. While wild-type cells were able to clear aggresomes almost completely, this effect was significantly impaired in Nrf1^KO^ cells (Fig 4A, 4B and 4C). These results suggested that Nrf1 may play a role in this selective form of autophagy and is required to clear aggresomes efficiently after proteasome inhibition.

**Figure 4.**
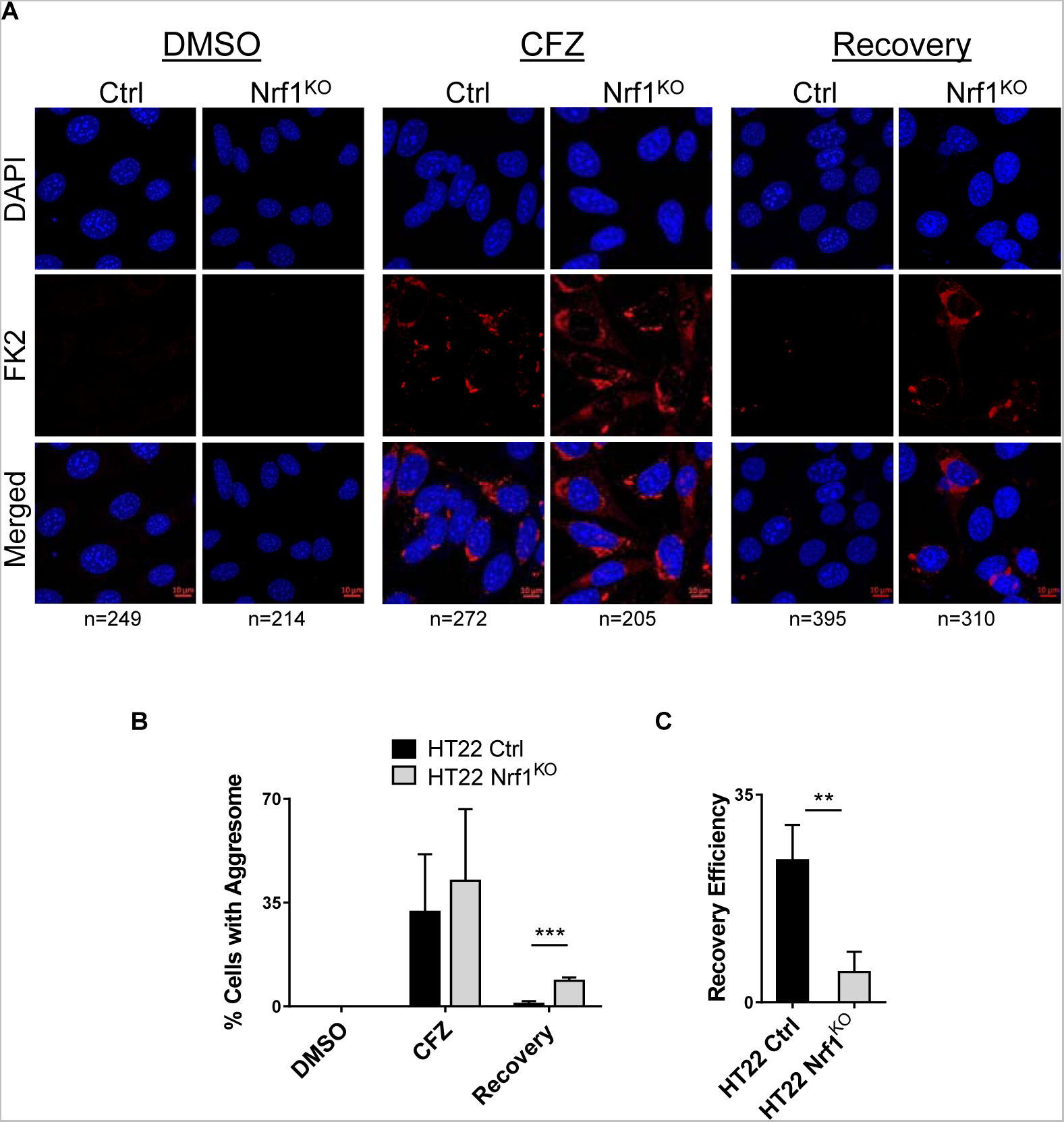
Nrf1 is required to clear aggresomes that are triggered by proteasome inhibition. **A.** HT22 wild-type (control; ctrl) and Nrf1^KO^ cells were treated with 50 nM CFZ for 20 h, then were washed and incubated with fresh media for another 20 h (Recovery period). Aggresomes were detected by confocal microscopy using FK2 stain. **B.** Percentage of cells with aggresomes under each condition (DMSO, CFZ and recovery) for both ctrl and Nrf1^KO^ cells are plotted, based on confocal microscopy analysis. Number of cells analyzed in each sample is noted underneath the images in (A). **C.** Recovery efficiency of ctrl and Nrf1^KO^ cells was calculated by dividing CFZ values by recovery values of ctrl and Nrf1^KO^ cells in (B). 3 biological replicates were used for confocal microscopy analysis. p-values were calculated by student’s t test. **,0.05, ***<0.005

### Deficiency of NGLY1, a positive upstream regulator of Nrf1, causes inhibition of compensatory autophagy and aggrephagy

N-linked glycosylation is a post-translational modification of Asparagine residues of proteins that are in the ER lumen (Suzuki et al., 2016). NGlycanase-1 (NGLY1) is an evolutionary conserved enzyme that removes N-linked glycosylation from cytosolic proteins (de- glycosylation). NGLY1 deficiency causes a rare congenital disorder characterized by developmental delay, seizures, neurological and liver malfunction among many other symptoms (Enns et al., 2014). On the molecular level, we and others have demonstrated that in this disease, deficiency of NGLY1 results in defective Nrf1 maturation due to the lack of its deglycosylation (Lehrbach et al., 2019; Lehrbach and Ruvkun, 2016; Tomlin et al., 2017). Given that de- glycosylation of Nrf1 is critical for its transcriptional activity, we asked if NGLY1 deficiency causes impairment of compensatory autophagy and aggrephagy in response to proteotoxic stress. To this end, we used Mouse Embryonic Fibroblast (MEF) cells that are wild-type, NGly1^KO^ and NGly1^KO^ stably expressing p110, the activated form of Nrf1 that does not require NGLY1 activity (NGly1^KO^ p110). First, to test if clearance of aggresomes is impaired in NGly1^KO^ cells, we treated wild-type, NGly1^KO^ and NGly1^KO^ p110 MEF cells with CFZ and let them recover in fresh complete media. Then, we stained the aggresomes with FK2 antibody and imaged them under the confocal microscope. As expected, all MEF cells induced aggresomes after proteasome inhibition (Fig 5A); however, wild-type cells cleared the aggresomes more efficiently than the NGly1^KO^ cells (Fig 5A and 5B). Importantly, addition of Nrf1 p110 in NGly1^KO^ cells resulted in the rescue of impaired aggresome clearance phenotype associated with NGly1 deficiency (Fig 5A and 5B), suggesting the requirement of Nrf1 in this process.

**Figure 5.**
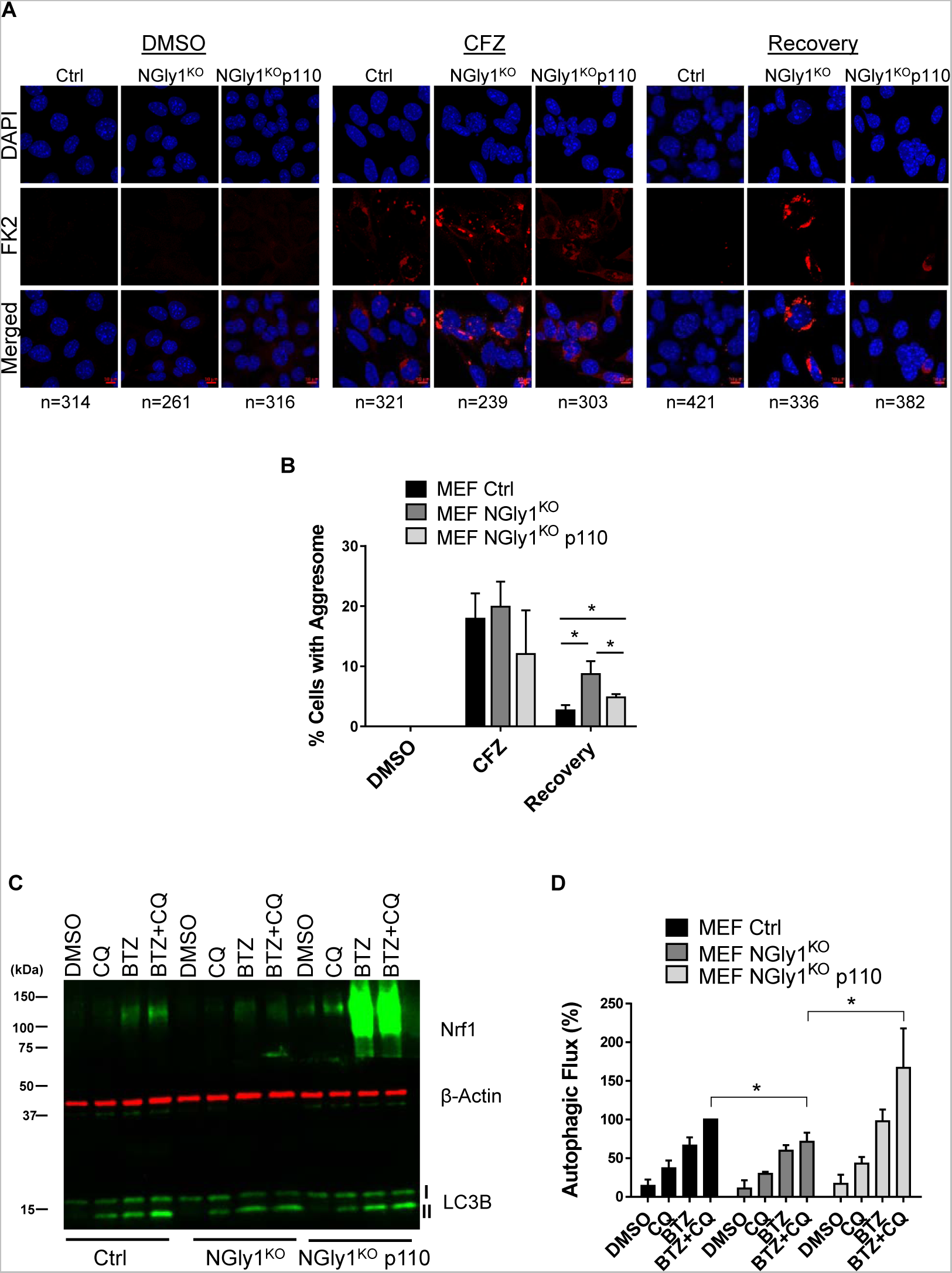
Deficiency of NGLY1 causes inhibition of compensatory autophagy and aggrephagy, which can be rescued by transcriptionally active Nrf1. **A.** Confocal images of FK2 labeled MEF ctrl, NGly1^KO^ and NGly1^KO^ p110 cells after treating them with DMSO or 25 nM CFZ for 20 h. Recovery samples were washed out after CFZ treatment and incubated in fresh complete media for 20h. **B.** Analysis of percentage of cells with aggresomes under each condition from (5A) are plotted, based on the confocal microscopy analysis. Number of cells analyzed in each sample is noted underneath the images in (B). **C.** Western blot analysis of LC3B, Nrf1 in MEF-ctrl, NGly1^KO^ and NGly1^KO^ cells expressing p110 construct treated with DMSO, 200 nM BTZ (20h), 60 μM CQ (3 h) or both BTZ (20 h) and CQ (3 h). β-Actin was used as a loading control. **D.** Representation of autophagic flux after normalizing LC3B-II levels to β- Actin levels from 5C. 3 biological replicates for qRT-PCR, microscopy and western blotting were used. p-values were calculated by student’s t test. *<0.5, **,0.05, ***<0.005

Next, we tested the autophagic flux in this disease model by western blotting for LC3B-II. Treatment of wild-type MEF cells with CQ and BTZ resulted in elevated LC3B-II levels, compared to DMSO (Fig 5C). As expected, treating cells with both CQ and BTZ caused an even higher LC3B-II level in wild-type cells, confirming that autophagic flux is activated in these cells by BTZ treatment. However, co-treatment of CQ and BTZ did not result an increase in LC3B-II levels in NGly1^KO^ cells (Fig 5C and 5D), compared to single treatments, suggesting that autophagic flux was impaired in NGly1^KO^ cells. Importantly, expression of Nrf1 p110 in NGly1^KO^ cells was able to rescue this impairment and restore autophagic flux, which was evident by elevated LC3B-II levels (Fig 5C and 5D). Overall, our data point to a critical role for Nrf1 in mediating proteasome inhibitor-induced compensatory autophagy and aggrephagy in this NGLY1 disease model.

## DISCUSSION

Our earlier work has firmly established a role for the transcription factor Nrf1 in responding to proteotoxic stress caused by inhibition of proteasome activity (Radhakrishnan et al., 2014; Radhakrishnan et al., 2010; Vangala and Radhakrishnan, 2019; Vangala et al., 2016). This response is characterized by transcriptional upregulation of proteasome genes resulting in the bounce-back of proteasome activity (Steffen et al., 2010). Our current study expands the role of Nrf1 in this context as a transcriptional regulator of ALP genes in addition to the proteasome genes. Thus, by increasing autophagic flux, Nrf1 provides the cells with an additional route to cope with proteotoxic stress. Proteasome inhibitors are known to trigger the formation of ubiquitylated protein aggregates (Kopito, 2000). Given that these aggregates are poor substrates of the proteasome, the ability of Nrf1 to induce autophagy provides the cells with the capacity to degrade these aggregates via aggrephagy as shown in our study.

Of note, a previous report from Goldberg and colleagues concluded that in response to proteasome inhibition, ALP genes p62/SQSTM1 and GABARAPL1 are rapidly induced to promote cell survival where p62 primarily acts by sequestering ubiquitylated proteins in inclusions (Sha et al., 2018). Although they showed that p62 is induced in an Nrf1 and NF-E2- dependent manner, the induction of GABARAPL1 and the other ALP genes together with activation of autophagy at a later time point was attributed to be Nrf1-independent. In contrast, our current study found a tight dependence of autophagy on the functional status of Nrf1. In Nrf1-depleted cells and in NGLY1-deficient cells (in which Nrf1 is inactive), using orthogonal methods, we found a clear defect in activating autophagy and the ability to clear aggresomes via aggrephagy. Furthermore, using GABARAPL1 as an example, we found direct binding of Nrf1 to its promoter region. Together with evidence from the ENCODE ChIP-seq datasets that indicate Nrf1 can bind to the regulatory regions of multiple ALP genes, our conclusions are consistent with a model in which Nrf1 can directly activate ALP genes to promote autophagy in the face of proteotoxic stress. While our manuscript was in preparation, a preprint article from Kobayashi and colleagues demonstrated that proteasome inhibitor mediated aggrephagy proceeded via direct activation of p62 and GABARAPL1 by Nrf1 (Hatanaka et al., 2023), thus further strengthening our model.

Besides providing mechanistic clarity, our findings also have important translational implications. In cancer therapies that utilize proteasome inhibitors (eg. Multiple myeloma), there is now a stronger justification to inhibit the Nrf1 pathway to increase the treatment efficacy. Although transcription factors such as Nrf1 remain undruggable directly, given the complexity of the pathway, there are ample opportunities to target activating enzymes such as the ATPase p97/VCP, N-glycanase NGLY1 and the protease DDI2, all of which are essential for the functional maturation of Nrf1 (Northrop et al., 2020). On the other hand, in certain neurodegenerative diseases (eg. Alzheimer’s disease) that exhibit ubiquitylated protein aggregates in the neurons, there could be interest in enhancing the Nrf1-dependent proteasome and autophagy pathways to clear the aggregates and improve proteostasis.

## MATERIALS AND METHODS

### Constructs

Nrf1-LentiCRISPRv2 was previously described (Vangala et al. 2016). To generate pLPCX-HA-Nrf1-3xFlag (a construct expressing p120 with N-terminal HA tag and C-terminal 3xFlag) we amplified Nrf1 ORF with HindIII/EcoRI-containing primers and cloned into pLPCXpuro. Similarly, we generated pLPCX-HA-p110Nrf1-3xFlag (a construct expressing p110, the active form of Nrf1) which is devoid of the N-terminal 104 amino acid of p120. These constructs are referred to as p120 and p110 respectively in this paper. pBABE-puro-mCherry-EGFP-LC3B was a gift from Jayanta Debnath (Addgene plasmid# 22418).

### Cell Culture

NIH-3T3, HT22, MDA-MB-231, MEF cells and their derivatives were grown in Dulbecco’s Modified Eagle Medium (DMEM) (Gibco) with 10% Fetal Bovine Serum (FBS, Atlanta Biologicals), 1X Penicillin-Streptomycin (Pen/Strep, Invitrogen) and 5 µg/mL Plasmocin Prophylactic (PP, InvivoGen). SH-SY5Y cells and its derivatives were grown in DMEM:Nutrient Mixture F12 with 10% FBS, 1X Pen/Strep and 5 µg/mL PP. All cells were maintained at 37°C in a humidified incubator with 5% CO_2_.

### Viral Transduction and Generation of Stable Cell Lines

To generate viral particles, 5.3 million HEK-293T cells were plated in 10-cm plates the day before transfection. Retroviral transfer plasmid (4µg), pUMVC (3.6µg) and pCMV-VSV-G (0.4µg) were transfected into the cells with Lipofectamine 3000 (Thermo Fisher, Cat No: L3000015), according to manufacturer’s instructions. To generate lentiviral particles, lentiviral transfer plasmid (4µg), pCAG-HIVgp (2µg), pHDM-G (1µg) and pCAG4-RTR2 (1µg) were used. Media supernatant containing viral particles was collected at 48 and 72 hr after transfection, then precipitated with PEG-it solution (Systems Bio). Concentrated viral particles were aliquoted and stored at -80°C.

The day before infection HT22 cells were plated in either 6-well plates or 10-cm plates at around 60% confluency. To generate stable cells for overexpression or for knock-out, one aliquot of viral particles with freshly added polybrene (4µg/mL) was used to infect cells plated in a 6- well plate, in serum-free media, which was replaced with complete media the next day. HT22 and NIH-3T3 cells that were infected with mCherry-GFP-LC3B virus were selected by FACS, two days after infection. Cells that were infected with Nrf1-LentiCRISPRv2 virus were selected by 1µg/mL puromycin for two weeks.

Nrf1-depleted cells in 10-cm plates were first infected by 2 aliquots of p120 viral particles. Three days after infection cells were split and infected again with 2 aliquots of viral particles for an efficient rescue. These cells were used to perform experiments after the second infection.

### Quantitative Reverse Transcription PCR (qRT-PCR)

Cells were plated in 6-well plates to reach 70% confluency at the time of treatment. Cells were treated with either 200 nM CFZ or 200 nM BTZ for 6 and 24 hr. At the end of treatment, cells were washed with PBS once and collected by scraping the cells in 1mL PBS. They were centrifuged at 14,000 rpm for 1 min at 4°C, PBS was aspirated off, and cell pellets were stored at -80°C. RNA isolation was performed by RNeasy Kit (Qiagen) with addition of DNAse (Qiagen) treatment. 1µg of RNA was used to make cDNA with iScript Reverse Transcriptase (Bio-Rad). qRT-PCR reactions were prepared with iTaq Universal SYBR green super mix (Bio-Rad) and ran in C1000 Touch Thermal Cycler (Bio-Rad). Depending on experimental conditions, either 18S or GAPDH was used as an internal control for normalization. Data were analyzed using CFX Manager 3.1 (Bio-Rad). The primers used for qRT-PCR are listed in Supplemental Table S2. Calculation of p-value was done by student’s t-test.

### Immunoblot Analysis

Cells were plated in 6-well plates to reach 70% confluency at the time of treatments. Cells were washed once with ice-cold PBS and then scraped in RIPA Buffer (20 mM Tris-HCl pH 7.4, 1% Sodium Deoxycholate, 1% Triton X-100, 0.1% SDS, 150 mM NaCl, 1 mM EDTA) containing protease and phosphatase Inhibitor cocktail (Fisher Scientific, Cat No:PI78447) on ice. Lysates were then sonicated at a low power setting for 5 seconds once, then incubated on ice for 20 min and centrifuged at 14,000 rpm for 20 min at 4°C. Protein concentrations were measured by Bradford assay. Samples for western blots were prepared with 4X Laemmli protein sample buffer (Bio-Rad, Cat No:1610747) with addition of 2-Mercaptoethanol on the day of lysis, followed by boiling for 7 minutes. 20 µg of protein was used for SDS-PAGE, and transferred on to Immobilon FL PVDF membranes (Fisher Scientific, Cat No:IPFL00005).

Membranes were blocked with Intercept TBS blocking buffer (Fisher Scientific, Cat No:NC1660550) for 1 hr, and then overnight incubation with specific primary antibodies, followed by incubation with secondary antibodies for 1 hr at room temperature. The antibodies used were, LC3B (Cell Signaling, Cat No:2775S) at 1:1000, β-Actin (Sigma-Aldrich, Cat No:A5441) at 1:10,000, Nrf1 (Cell Signaling, Cat No:8052S) at 1:1000 and GABARAPL1 (Cell Signaling, Cat No:26632S) at 1:1000. IRDye 800CW Goat-anti-Rabbit antibody (VWR, Cat No:102673-330) and IRDye 680RD Goat-anti-Mouse antibody (VWR, Cat No:102673-408) at 1:20,000 dilutions. After secondary antibody incubation, membranes were washed with TBST (4 times, 5 minutes each) and dried for 1 hour in the dark before imaging with LiCOR Odyssey Fc Imaging System. Quantification of western blot signals was done by Image Studio Lite V5.2 software, and student’s t-test was used to determine p-values. All graphs were plotted by using GraphPad Prism software.

### LC3B Puncta Analysis

0.25 million HT22 mCherry-GFP-LC3B cells were plated in 6-well plates with cover slips. Cells were treated with either DMSO or 200 nM BTZ for 18 hr. After treatment cells were washed with PBS 1X and fixed with ice-cold methanol for 10 min, followed by PBS wash 3X and mounted on to slides with Prolong Gold Antifade Mountant with DAPI. Images were captured using spinning disc confocal microscopy with 63X objective. The GFP and RFP (mCherry) signals were imaged using a line sequential scan setting with excitation laser lines at 488 and 543 nm, respectively. The emission signals were collected at 495 to 530 nm (GFP, channel 1) and 590 to 650 nm (RFP, channel 2). Puncta staining was quantified by ImageJ software, with the help of ‘Green and Red Puncta Colocalization’ macro, developed by Daniel J. Shiwarski, Ruben K Dagda and Charleen T. Chu. NIH-3T3 mCherry-GFP-LC3B cells were plated in glass-bottom 96-well plates, and after fixation, images were acquired using high-content microscope (Operetta, PerkinElmer). Nine fields per well were captured and the images were analyzed using the manufacturer’s software.

### Aggresome Assay

0.25 Million HT22 and 0.4 million MEF cells were treated with either 50 nM CFZ or with DMSO for 20 hr on cover slips in 6 well plate. The cells that received a recovery period after a 20hr CFZ treatment were washed with PBS twice and then supplied with fresh complete media for 20 hr. At the end of treatment, cells were washed with PBS, fixed with ice-cold methanol, blocked with blocking solution (1% BSA and 0.3% Triton-X in PBS) and stained with FK2 antibody at a 1:500 dilution (Cayman Chemicals; Milipore) overnight (Klickstein et al., 2020; Mukkavalli et al., 2021) for overnight, followed by washing and incubation with Goat-anti-Mouse Alexa-flour 555 (Invitrogen) for 1 hr. After incubation, cells were washed once in blocking solution and then incubated for 5 min in Hoechst diluted 1:10,000 in PBS. Hoechst was aspirated off followed by one wash of PBS and two of deionized water before cover slips were mounted. Spinning disc confocal microscopy with 63X objective was used to image the cells. ImageJ with Aggrecount macro (Klickstein et al., 2020) was used to quantify images.

### Chromatin Immunoprecipitation (ChIP)

ChIP assay was performed as described previously (Vangala and Radhakrishnan, 2019). Briefly, cells were cross linked for 10 min with 1% (w/v) formaldehyde, followed by quenching with 0.125M glycine. After cold pbs washes 2X, pellets were collected at 800 × g at 4 °C. Chromatin was isolated using EZ-Magna ChIP A/G (Millipore) kit according to the manufacturer’s protocol. Covaris M220 was used with 10% duty factor (df) for 12 min for chromatin shearing. Supernatant was collected from sheared chromatin at 10,000 × g for 10 min and precleared with 20 μl of protein A/G magnetic beads for 1 hr at 4 °C. Immunoprecipitation was completed with 5 μg of specific antibody for 50 μl of precleared chromatin overnight at 4 °C. After immunoprecipitation, beads were consequently washed with a series of buffers and eluted in elution buffer according to the manufacturer’s protocol. Qiagen columns were used for purification of eluants. qPCR was used to analyze the chromatin pulldown using specific primers shown in Supplemental Table S2.

## ACKNOWLEDGEMENTS

We thank Dr. Malavika Raman (Tufts University) and her laboratory for advice on aggresome clearance assays. We thank Dr. Tadashi Suzuki (RIKEN) for NGly1-knockout MEFs and Dr. Lars Steinmetz (Stanford University) for SH-SY5Y-Nrf1-CRISPRi cells. S.K.R was supported by an NIH award R01GM132396 and pilot awards from Grace Science Foundation and Albert Einstein College of Medicine Nathan Shock Center. H.A.B was supported by a diversity supplement to the NIH award R01GM132396. Services in support of the research project were generated by the VCU Massey Cancer Center Microscopy Shared Resource, supported, in part, with funding from NIH-NCI Cancer Center Support Grant P30 CA016059.

---

**Supplemental Figure S1.**
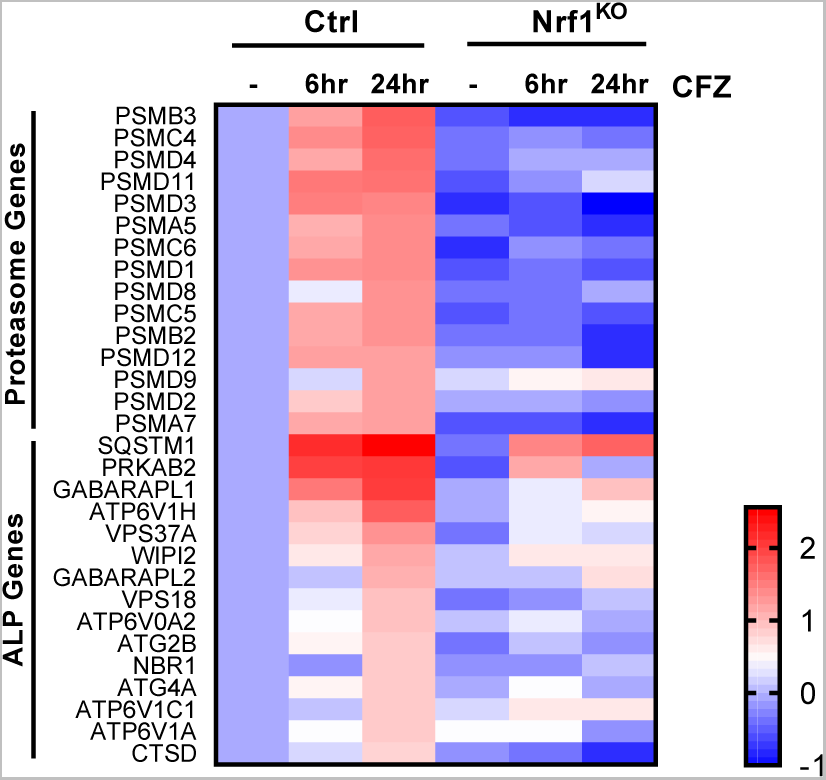
Nrf1-dependent regulation of proteasome and autophagy lysosomal pathway (ALP) genes in response to carfilzomib (CFZ). Heatmap analysis of RNA-seq data (GSE144817) obtained from wild-type (control; ctrl) and Nrf1^KO^ NIH-3T3 cells treated with DMSO or with 200 nM CFZ for 6 h or 24 h. Log2 fold changes are shown.

**Supplemental Figure S2.**
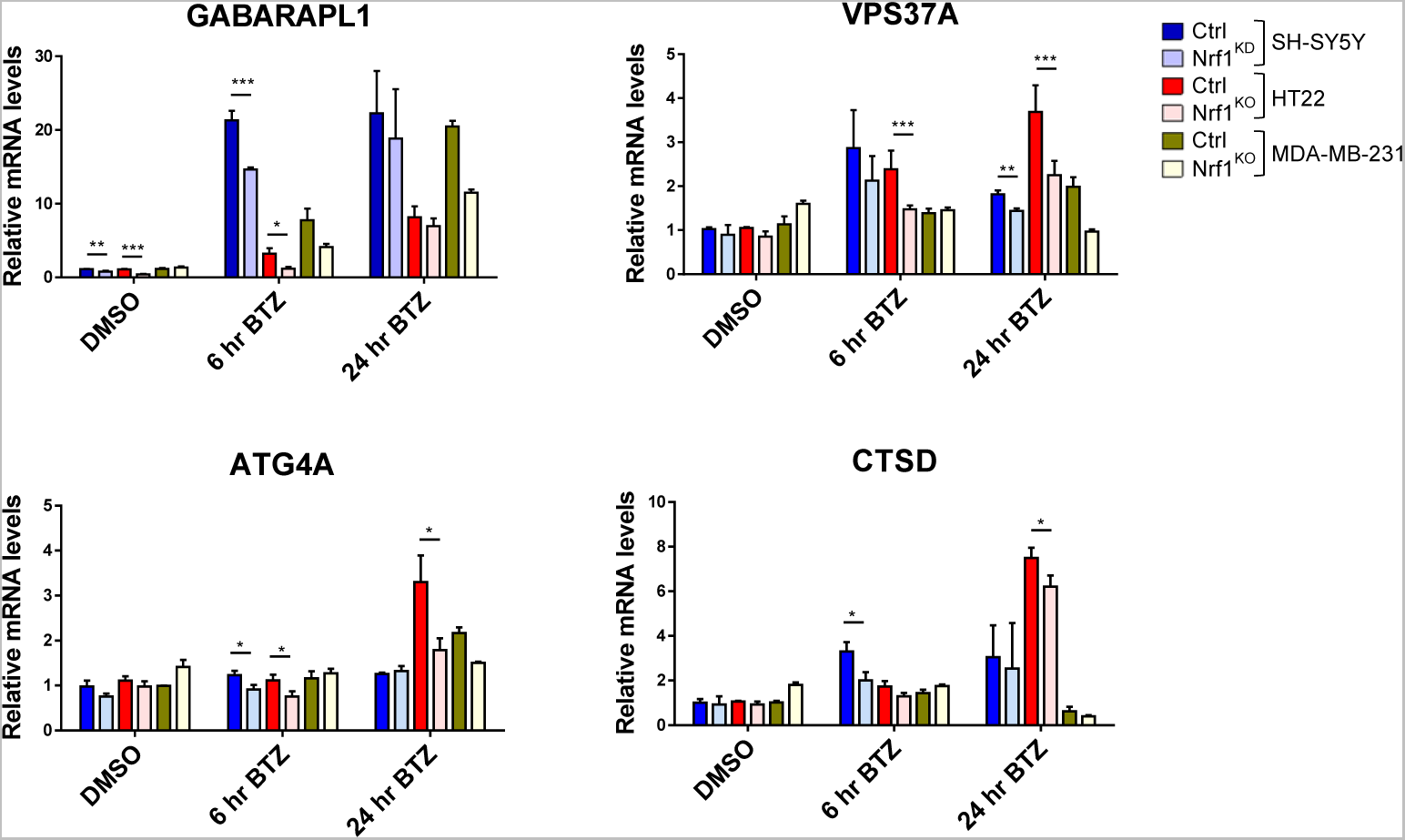
Nrf1 regulates expression of Autophagy-Lysosome Pathway (ALP) genes upon proteasome inhibition. qRT-PCR analysis of SH-SY5Y, HT22 and MDA-MB-231 cells that are control (Ctrl) or Nrf1-depleted (KO or KD) were treated with either DMSO or 200 nM bortezomib (BTZ) for 6 h or 24 h. Expression levels of GABARAPL1, VPS37A, ATG4A and CTSD were analyzed using gene specific primers as shown. 18s rRNA or GAPDH levels were used for normalization.

**Supplemental Figure S3.**
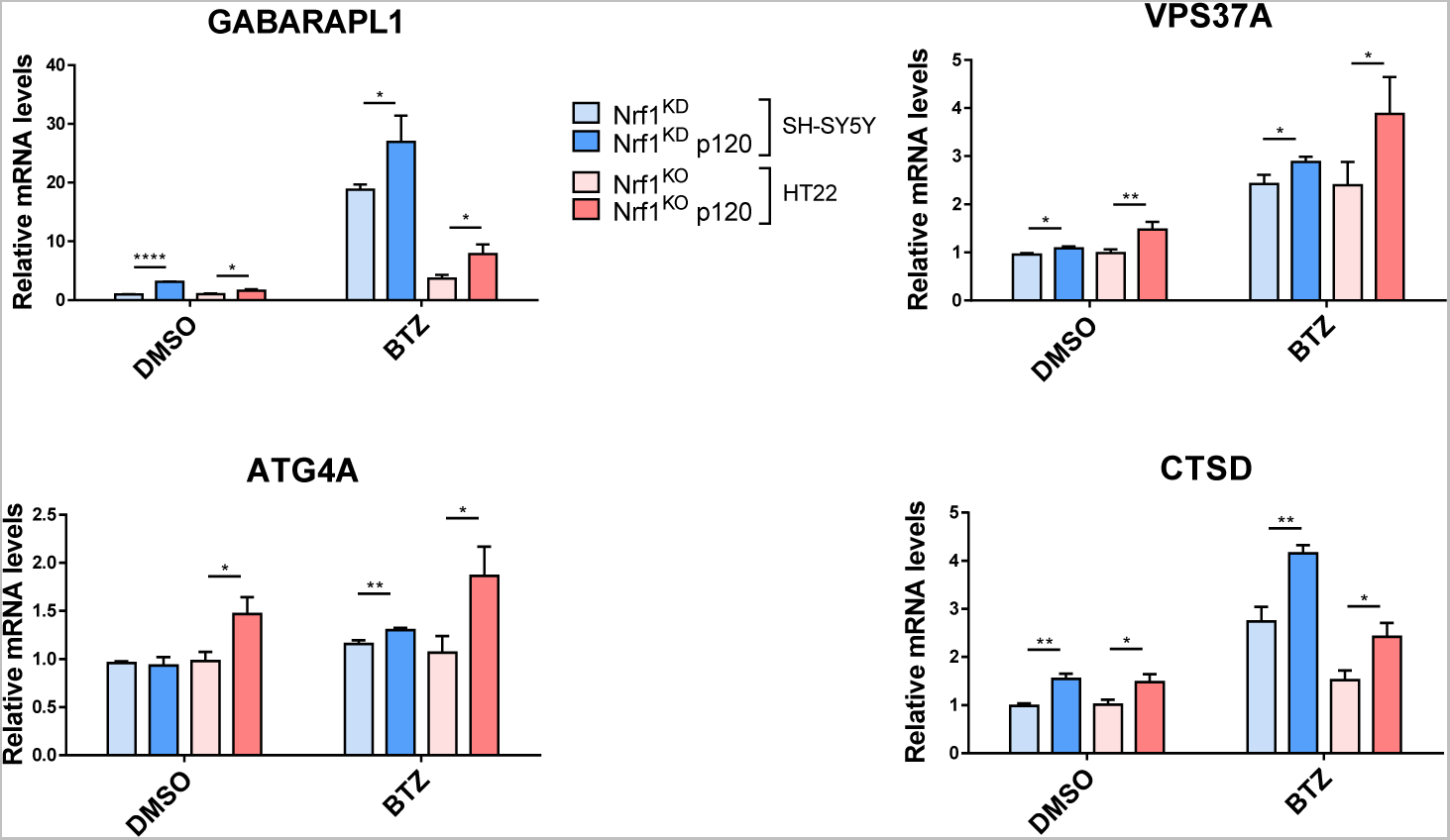
Adding back Nrf1 in Nrf1-deficient cells rescues suppressed expression of ALP genes upon proteasome inhibition. SH-SY5Y-Nrf1^KD^ and HT22-Nrf1^KO^ cells were infected with Nrf1(p120). Both SH-SY5Y-Nrf1^KD^, p120 rescue and HT22-Nrf1 ^KO^, p120 rescue cells were treated with 200 nM BTZ for 6 h and then analyzed by qRT-PCR to measure the expression levels of indicated genes and mRNA levels of 18s rRNA levels was used for normalization.

**Supplemental Figure S4.**
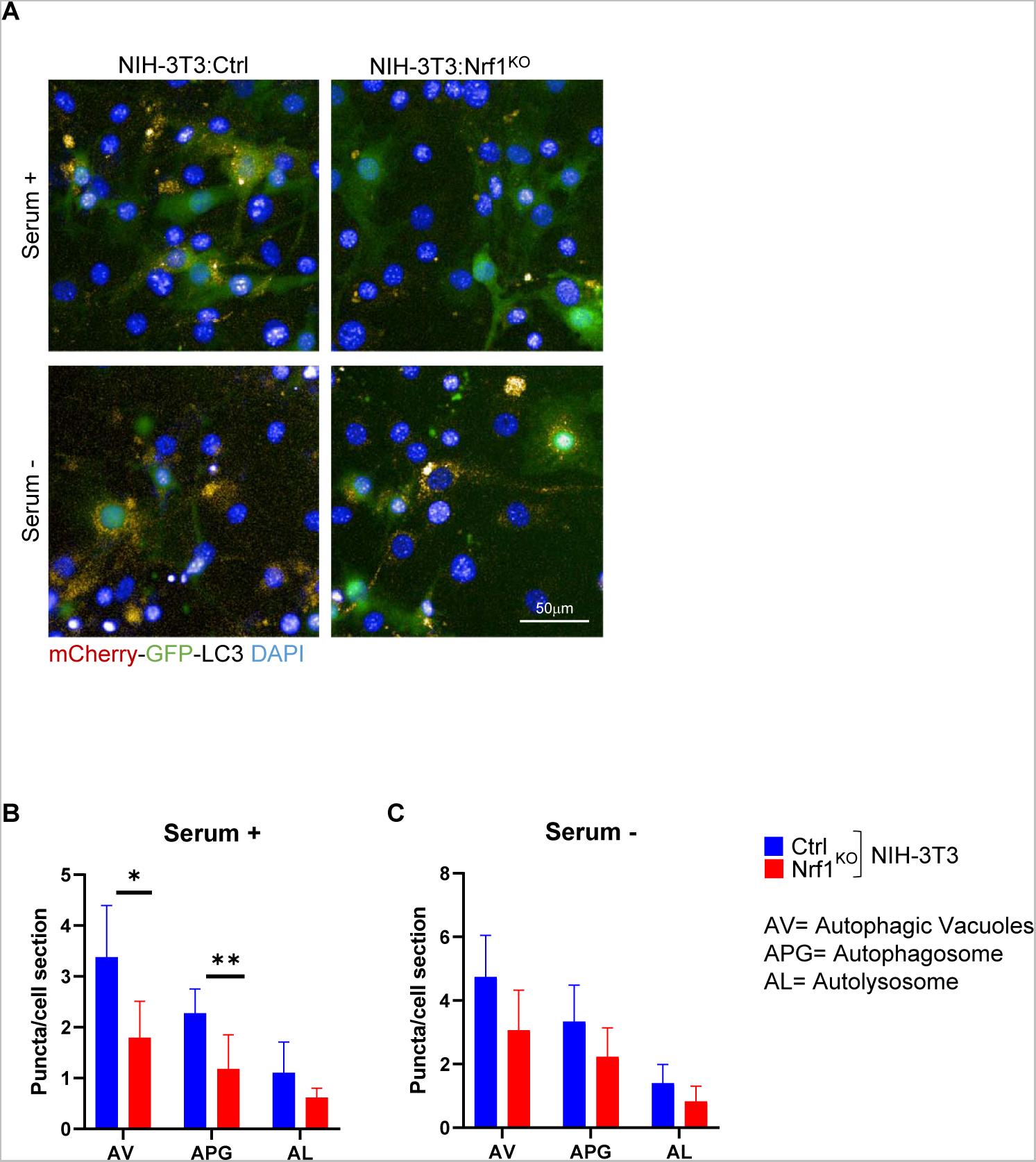
Nrf1^KO^ cells display defects in basal and starvation-induced autophagy. NIH-3T3 cells that are wild-type control (ctrl) or Nrf1^KO^ expressing the tandem reporter mcherry-GFP-LC3 were incubated in serum supplemented (serum +) or serum deprived (serum -) media for 6 h. Number of red puncta (autophagic vacuoles; AV), yellow puncta (autophagosomes; APG) and red only puncta (autolysosomes; AL) were quantified as the average number of fluorescent puncta per cell. Representative images are shown in **A** and the quantification of puncta are shown in **B** and **C**. Differences were significant for *p<0.05 and **p<0.001.

**Supplemental Table S1.** Nrf1 binding locations derived from ENCODE project database (Identifier: ENCSR245QUM)

| **Autophagy / Lysosomal Genes** | **Nrf1 binding peak's approx. distance from TSS (+1)** |
| --- | --- |
| ATG4A | -817, -349 |
| ATP6V0A2 | -256 |
| ATP6V1A | +96 |
| CTSD | -98, +638 |
| GABARAPL1 | -444, +192 |
| GABARAPL2 | +102, +472 |
| PRKAB2 | +102, +811 |
| SQSTM1/p62 | -1810, -1360, -326, +92 |
| VPS18 | -87, +587 |
| VPS37A | -67, +603 |
| WIPI2 | -86 |

**Supplemental Table S2.**
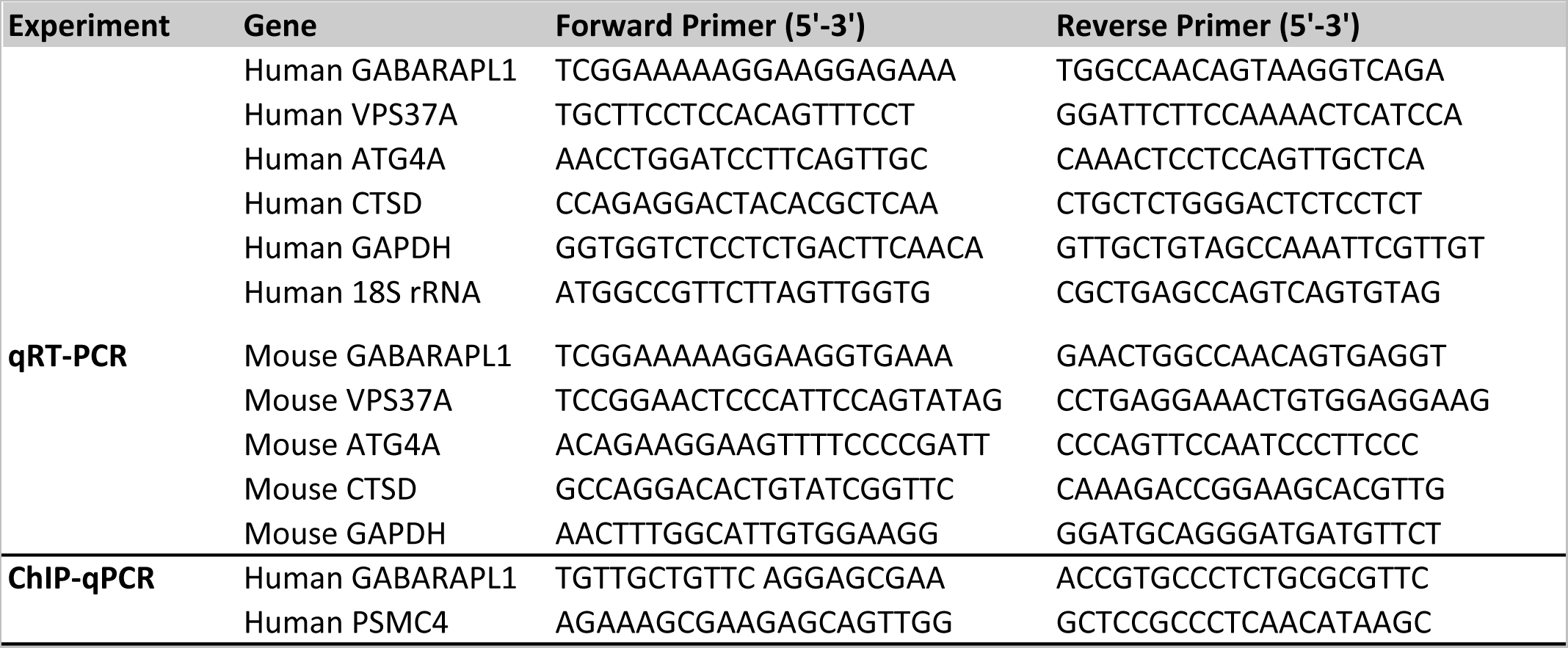
Primers used in quantitative RT-PCR (qRT-PCR) and chromatin immunoprecipitation (ChIP)-qPCR assays.

